# CycloCross: Adversarial single-cell RNA-seq data translation across species

**DOI:** 10.64898/2026.09.24.754263

**Authors:** Olympio Hacquard, Yuya Tokuta, Yusuke Imoto, Tomonori Nakamura, Shuto Deguchi, Masahiro Nagano, Mitinori Saitou, Yasuaki Hiraoka

**Affiliations:** Institute for the Advanced Study of Human Biology, Kyoto University, Yoshida-Konoe-Cho, Sakyo-Ku, Kyoto, 606-8501, Japan; Department of Anatomy and Cell Biology, Kyoto University, Yoshida-Konoe-Cho, Sakyo-Ku, Kyoto, 606-8501,Japan; Hakubi Center, Kyoto University, Yoshida-Konoe-Cho, Sakyo-Ku, Kyoto, 606-8501, Japan; Department of Biological Engineering, Massachusetts Institute of Technology, Cambridge, MA 02139, USA; Center for iPS Cell Research and Application (CiRA), Kyoto University, 53 Kawahara-cho, Shogoin, Sakyo-ku Kyoto, 606-8507, Japan; Inamori Research Institute for Science, Inamori Foundation, 620 Suiginya-cho, Shimogyo-ku Kyoto, 600-8411, Japan; Center for Advanced Intelligence Project, RIKEN, 1-4-1 Nihonbashi, Chuo-ku Tokyo, 103-0027, Japan

**Keywords:** scRNA-seq, CycleGAN, cross-species analysis, data translation, adversarial networks, batch correction, contrastive genes

## Abstract

Single-cell RNA sequencing (scRNA-seq) has transformed our understanding of cellular heterogeneity, yet most datasets are limited to a handful of model organisms, leaving critical gaps in cross-species biology. Existing computational methods for integrating multi-species scRNA-seq data often rely on one-to-one ortholog mapping, which fails to account for gene duplications, losses, or functional divergences. To address these challenges, we introduce CycloCross, an adversarial method based on the CycleGAN architecture, designed to translate scRNA-seq data between species without requiring a priori gene correspondences. Unlike generative models that synthesize data from noise, CycloCross leverages real data from a source species, enabling more realistic and data-efficient translations. We demonstrate its effectiveness in germ cell development, translating data between mouse, macaque, and human with conserved yet divergent transcriptional programs. CycloCross not only preserves temporal dynamics but also enables interpolation and extrapolation of gene expression at missing time points, offering predictions for experimentally inaccessible states. Furthermore, CycloCross generates biologically plausible samples even with limited data, outperforming traditional generative models.

## Introduction

The advent of single-cell RNA sequencing (scRNA-seq) has revolutionized our ability to dissect cellular heterogeneity and uncover the molecular underpinnings of complex biological systems. However, the vast majority of scRNA-seq datasets are generated from a limited number of model organisms, leaving significant gaps in our understanding of species-specific biology. Bridging these gaps requires innovative computational approaches capable of translating scRNA-seq data across species, thereby enabling the generation of high-resolution transcriptional atlases for uncharted organisms.

Unifying data from multiple species presents a challenge in removing species-specific heterogeneity. This problem inherently involves aspects of batch-effect correction, for which many methods have been developed [11, 16, 25, 42]. Some methods specifically address the multi-species aspect [44, 48, 60]. Inferring gene correspondence is a challenge, for which several methods have been proposed, e.g. SAMap [48] that refines many-to-many homologous gene relationships, Species-OT [49] that uses optimal transports to find gene correspondences between species and is purely expression-based, SATURN [44] and UNIFY [58] that both use protein large language models. While this latter approach expands the scope of cross-species integration, it still relies on precomputed functional annotations, which may introduce biases, particularly for less well-characterized species. Moreover, inferring gene correspondence is not sufficient to translate gene expressions across species. A strong advantage of the method developed in this paper is that we develop an integration method completely agnostic to gene matching and gene annotation.

Batch integration methods typically modify low-dimensional representations of gene expression to enable joint analysis of datasets from different species. Developing a method that translates data from one species to another, in addition to removing species-specific batch effects, is motivated by two key applications. First, it allows the transfer of experimental results from a model species to a species of interest when the experiment can only be conducted on the model species for practical, economical, or ethical reasons. This has been explored in previous work, such as the Icebear method that uses a variational autoencoder (VAE) to transform scRNA-seq data from one species into the style of another [55]. However, this method remains restricted to one-to-one ortholog mapping and does not assess the realism of the generated cell profiles, a critical consideration for downstream biological interpretations. Second, it enables in silico data generation for a given species using real data from another species. This is particularly valuable in biology, where analyses are often limited by small sample sizes [5, 37]. Augmenting real data with synthetic data has been shown to improve the robustness of analyses, both theoretically and in practice [32, 43]. In the context of scRNA-seq data, numerous neural network architectures have been introduced to generate in silico data, including VAEs [13, 31, 40], Generative Adversarial Networks (GANs) [34, 61], diffusion models [28], and large language models [30]. All these data generation methods ultimately learn to generate in silico data by randomly sampling from a latent space and transforming the sampled vector into a gene expression profile. In contrast, a method based on data translation generates new synthetic data from real data of another species. If the two species are sufficiently similar, the translation required to map a gene expression profile to a different species is much simpler than the transformation needed to map a random vector of noise to a gene expression profile. Consequently, a method based on translating data from existing species is expected to provide more realistic samples while requiring less data.

We introduce CycloCross, an adversarial method to translate single-cell RNA-seq data between species. CycloCross relies on the CycleGAN architecture [59] to translate single-cell RNA seq data between different organisms. CycleGANs have been previously applied to biological data to combine different modalities such as scRNA-seq and ATAC-seq data [52], to integrate spatial information with scRNA-seq data [2] and to enhance the resolution of single-nucleus RNA-seq data to match that of scRNA-seq [8]. CycloCross is a symmetric method that takes two scRNA-seq datasets from two different species and translates each in the style of the other species. It employs an adversarial approach by simultaneously training four neural networks: two generators, each of which takes a cell from a given species and translates it into a cell of another species, and two discriminators that ensure the generated cells are indistinguishable from real cells. The generators can take vectors of any dimension as input and output, allowing CycloCross to operate on species with different gene sets. We demonstrate that CycloCross generates realistic samples. In particular, we investigate CycloCross in the context of germ cell development, translating data between mouse, macaque, and human. Germ cell development in mammalian closely related species is governed by a well-conserved transcriptional network, and species differences between mice and humans have also been extensively investigated through wet-lab experiments [23, 24]. We show that CycloCross respects the time-evolution of this data and can be used for gene expression interpolation or extrapolation at missing time points, thereby enabling predictions of transcriptional states that are difficult to observe experimentally. Finally, we demonstrate that CycloCross can also be used to analyze which genes exhibit species-specific expression patterns, offering insights into the molecular mechanisms underlying evolutionary divergence and bringing us a step closer to answering the question of what makes us human.

## Results

### CycloCross translates scRNA-seq data across species: method overview

CycloCross takes as input scRNA-seq data from two species, *S*_1_ and *S*_2_, represented as gene-by-cell expression matrices. Following a preprocessing pipeline (described in Methods) independent for each species, we end up with *p*_1_ genes for *S*_1_ and *p*_2_ genes for *S*_2_. We do not assume any ortholog mapping between the genes of each species. In particular, *p*_1_ and *p*_2_, as well as the corresponding gene lists, may be different. CycloCross translates cells from *S*_1_ into the expression space of *S*_2_, and vice versa (Fig. 1a). We refer to the translated datasets as *S*_2_-like *S*_1_ and *S*_1_-like *S*_2_, respectively.

**Figure 1:**
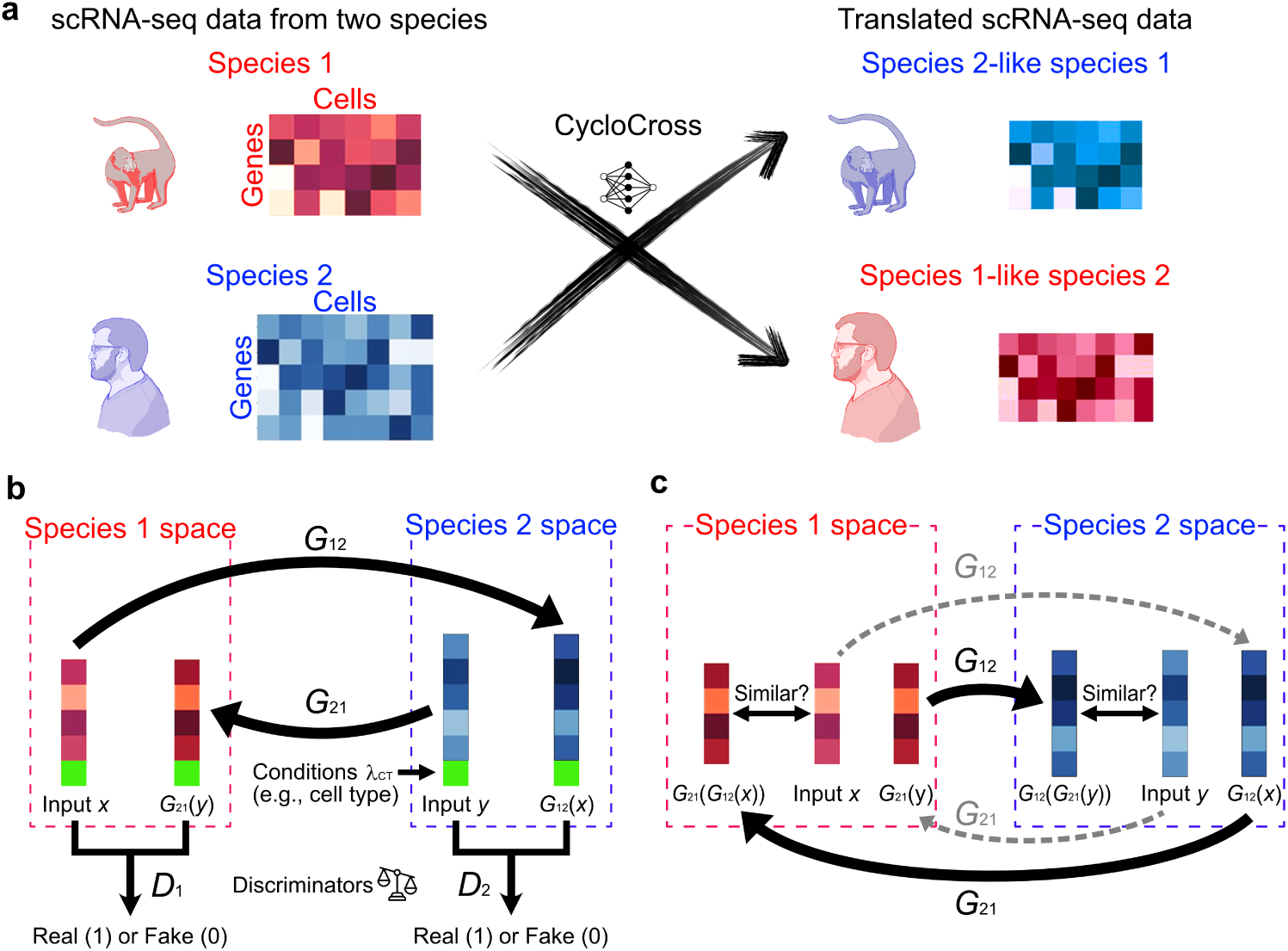
CycloCross pipeline. **a**, General principle, CycloCross takes scRNA-seq data from two species and translates each data in the style of the other species. **b** Adversarial data generation. A condition vector is appened to the input scRNA-seq data. Generators translate data across species while discriminators classify between real and simulated data. **c** Cycle principle: the translation of translated data back to the space of the original species must resemble the original data.

The translation procedure is based on the CycleGAN approach [59], initially developed for images, which takes unpaired data from two distributions and generates data from one distribution *in the style* of the other distribution. This is done through an adversarial approach with four distinct neural networks: a generator *G*_12_ : R*^p^*^1^ *→* R*^p^*^2^ to map cells from *S*_1_ to cells from *S*_2_. Another generator *G*_21_ : R*^p^*^2^ *→* R*^p^*^1^ to map cells from *S*_2_ to cells from *S*_1_. A discriminator *D*_1_ : R*^p^*^1^ *→* [0, 1] to ensure that cells generated in the style of *S*_1_ are realistic. Another discriminator *D*_2_ : R*^p^*^2^ *→* [0, 1] to ensure that cells generated in the style of *S*_2_ are realistic.

Similar to the GAN [12] and Wasserstein-GAN (WGAN) [4] frameworks, CycloCross generates data in an adversarial way. Discriminators are binary classifiers that aim to distinguish between fake and real data, where generators try to fool discriminators by creating data as realistic as possible. Discriminators are sometimes referred to as "critics" in the case of WGAN. Unlike the GAN approach, where generators take a random vector from a latent space as input, the generators in CycloCross are fed with single-cell RNA seq data from another species. Due to evolutionary similarities between the two species, the generating task is thus facilitated as demonstrated later.

Denote by 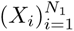 the filtered scRNA-seq data from species *S*_1_ constituted of the gene expression of *N*_1_ cells, and by 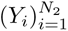 those of species *S*_2_, constituted of *N*_2_ cells. We assume that the scRNA-seq data 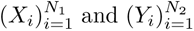 are respectively distributed according to two distributions P*_X_* and P*_Y_* . The classical GAN loss for generating data from *S*_2_ is:

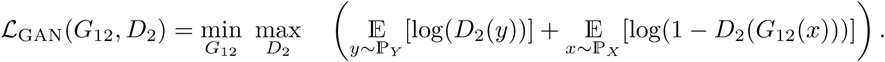

The GAN loss to generate data from *S*_1_ in the style of *S*_2_ is similar. This loss illustrates the competition between *G*_12_ and *D*_2_. The discriminator *D*_2_ tries to maximize this quantity by classifying real data sampled from P*_Y_* as 1 and fake data from P*_X_* as 0. The generator *G*_12_ on the other hand, tries to fool the discriminator by forcing it to classify the data generated from P*_X_* as 1 (Fig. 1b). Along the data pass, a mis-classification by the discriminators decreases the value of the loss function, while a correct classification increases it.

To ensure that the generated cells correspond to translations of existing cells in the other species (and not simply realistic cells), we add a cycle loss [59] (Fig. 1c). This loss ensures that translating a cell into the target species and then back to the original species must be close to the original cell:

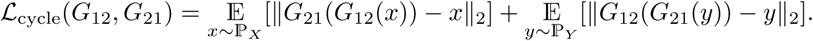

The total loss combines the GAN loss and the cycle loss:

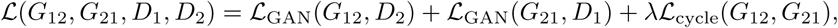

where *λ* is a real parameter balancing between the two objectives (generating realistic cells and providing accurate translations). Implementation details regarding the networks and the list of hyperparameters are provided in Methods.

To take into account any meta information we have about the data, such as tissue type, cell type or time information, we consider conditional adversarial networks [36]. In practice, this is done by concatenating the input vectors with a *condition vector c*. This is a common strategy in generative models for scRNA-seq data [28, 55]. Depending on the nature of *c*, several strategies can be derived: when *c* is a discrete information such as the cell type, we consider a one-hot encoding of the cell type information. When *c* is a continuous information such as a real-valued time stamp, we simply concatenate it at the end of the vector of gene expression. This is the case of the germ cell development data that we study extensively in this paper. Note that we do not require matching time stamps between *S*_1_ and *S*_2_. In both cases, the importance of this information is calibrated with a parameter *λ*_CT_ prior to concatenation to the vector of gene expressions (Fig. 1b).

### CycloCross translates temporal data with different time stamps between species

The main strength of CycloCross is its translation ability by transforming every cell of *S*_1_ as if they had been sampled from *S*_2_. Here, we demonstrate the ability of CycloCross to efficiently map cells of a species towards cells of another species while respecting the temporal organisation of cells. We use CycloCross to do every pairwise translation between macaque, mouse and human data. All these data consist of primordial germ cell-like cells (PGCLCs) induction systems from stem cells observed at different time stamps and different stem cell states. More precisely, human data consist of embryonic stem cells (ESCs), incipient mesoderm-like cells (iMeLCs), and PGCLC data measured after one, two, three and four days. Macaque data consist of ESC and PGCLC measured after one, two, four, six and eight days. Mouse data consist of epiblast-like cells (EpiLCs) and cells from days 2, 4, and 6 after the initiation of PGCLC induction from EpiLCs. Human data comes from a different lab than macaque and mouse data, demonstrating the ability of CycloCross to translate data between species, even when additional experimental batch effects are present. The three datasets differ substantially in sample size: 11,277 cells for human, 8,432 cells for macaque, and only 1,761 cells for mouse. This imbalance is particularly pronounced in the human-mouse comparison, where the mouse dataset contains far fewer cells and lacks several time points available in the human dataset (see Methods section for details).

CycloCross maps each cell of *S*_1_ measured at time *t* to a cell of *S*_2_ measured at time *t*, even if the scRNA-seq data from *S*_2_ contains no measurement at time *t*. For each pair of species, we embed original cells from a species and fake cells from the other species in the same UMAP space, where we separately plot the true cells and the counterfeit cells (Figs. 2a and 2b). Thanks to its time-conditional approach, CycloCross preserves the temporal organization of the translated cells, such that cells with similar time stamps remain clustered together in the target species space and translated cells are placed near real cells with related developmental stages. Notably, this behavior is also observed in the human–mouse translation despite the strong imbalance in cell numbers between the human and mouse datasets, suggesting that CycloCross can learn temporally consistent translations even when the source species is represented by substantially fewer cells.

**Figure 2:**
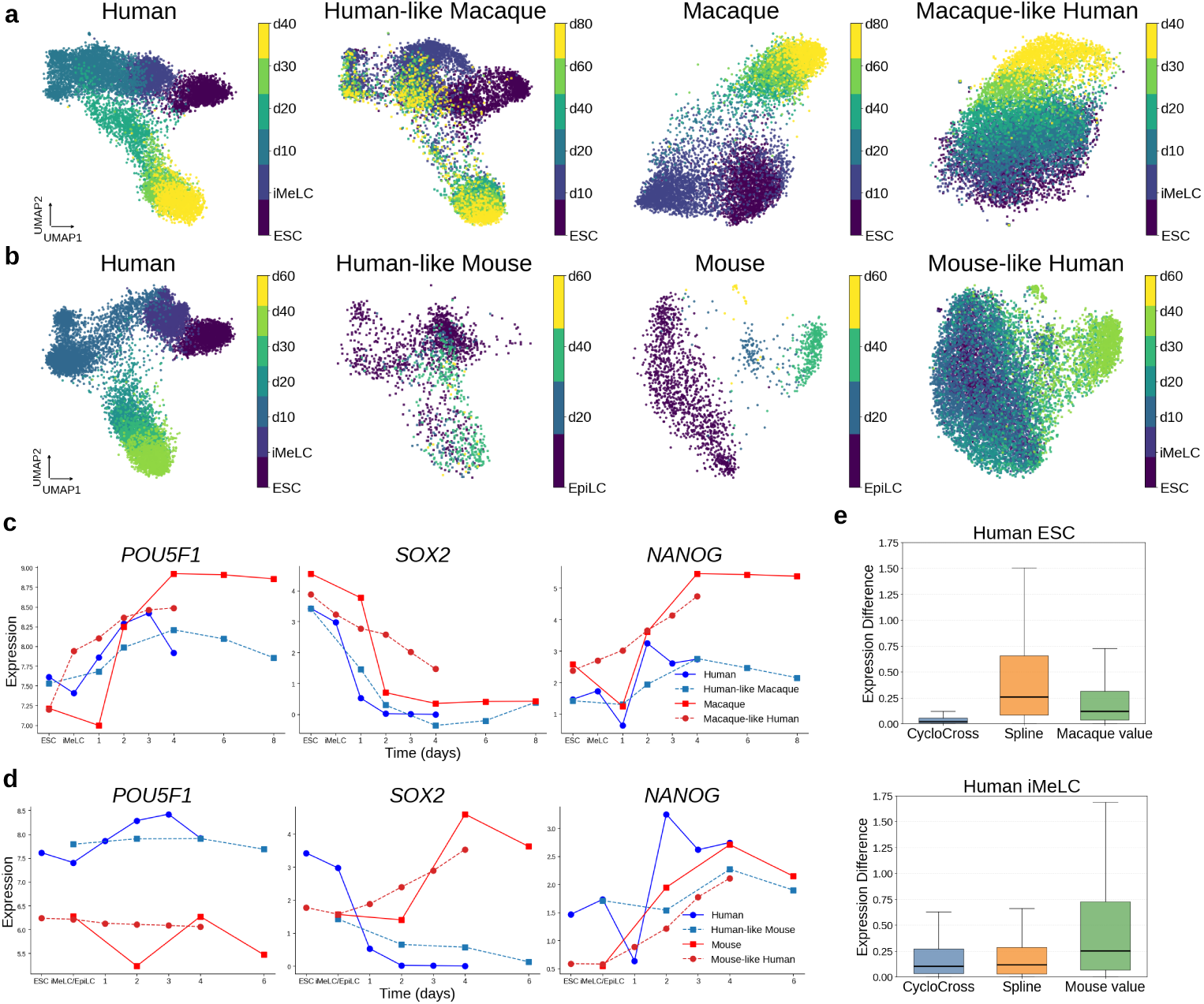
Translation results and verifications of CycloCross for scRNA-seq datasets from three-species’s PGCLC induction system with different time points. **a**, UMAP plots of human and macaque data translated by CycloCross, colored by time stamp. Real and simulated data are embedded in the same UMAP space, and then plotted separately. **b**, UMAP plots of human and mouse data translated by CycloCross, colored by time stamp. Real and simulated data are embedded in the same UMAP space, and then plotted separately. **c**, Evolution of the gene expression of *POU5F1*, *SOX2* and *NANOG* for Human-Macaque translated data. **d**, Evolution of the gene expression of *POU5F1*, *SOX2* and *NANOG* for Human-Mouse translated data. **e**, Box plot of error for imputation of pseudo-bulk gene expression of Human ESC from ESC macaque data and of human iMeLC from mouse EpiLC data.

To assess whether CycloCross captures both the temporal dynamics and the magnitude of gene expression across species, we compute the average log-normalized expression of each gene across cells at each time point for both the real scRNA-seq data and the CycloCross-translated data. This allows us to compare the temporal trajectories of gene expression between real and translated datasets across species pairs, overcoming the difficulty in aligning the cell types across species. We specifically examine three key transcription factors: *POU5F1*, *NANOG*, and *SOX2*, which are crucial for pluripotency and known to exhibit species-specific expression levels or dynamic temporal changes during PGCLC induction [23, 45] (Figs. 2c and 2d). *POU5F1* exhibits an increasing trend in both macaque and human datasets, with CycloCross accurately recapitulating this pattern. Notably, a decrease in expression at the end of the available data is observed in the human data, a feature also faithfully reproduced by CycloCross. When translating between mouse and human data, CycloCross preserves the overall stability of *POU5F1* while adjusting its baseline expression toward the target species’ level. For *SOX2*, both human and macaque datasets display a decreasing trend, which CycloCross effectively captures. In contrast, the mouse dataset exhibited a strong increase in expression, peaking at day 4. When trained to translate between human and mouse data, CycloCross successfully reconstructed these opposing trends, demonstrating its ability to preserve species-specific expression patterns. Lastly, for *NANOG*, both macaque and human datasets showed a sharp minimum at day 1, followed by an increase to high expression levels. CycloCross approximates these global trends and expression levels appropriately, though it fails to capture the initial decrease in macaque data and only partially reproduces it in human data. For the mouse dataset, where no such decrease is observed and gene expression increases monotonically until day 4, CycloCross again accurately captures the trend.

We evaluate the ability of CycloCross to predict bulk-level gene expression in a target species using data from a source species, focusing on two experimental setups. First we infer human embryonic stem cell (ESC) gene expression using macaque ESC data. CycloCross is benchmarked against two approaches: a cubic spline extrapolation trained on the remaining human time-series data, and a naive imputation using macaque ESC expression values directly. This evaluation mirrors a previously established methodology [55]. Across 1,144 transcription factors with one-to-one orthologs between human and macaque, CycloCross significantly outperforms both methods, and does not present any aberrant expression value (Fig. 2e). This demonstrates the ability of CycloCross to predict missing cell states using information from a source species. Next, we predict human intermediate mesoderm-like cell (iMeLC) expression using mouse epiblast-like cell (EpiLC) data as it is the closest available state. CycloCross is compared to cubic spline interpolation and naive imputation of mouse EpiLC values. Averaging across 742 orthologous transcription factors, CycloCross performs only slightly better than spline interpolation. Here, splines are particularly adapted to model monotonic transitions between ESC and early PGCLC stages. This result reflects the unique challenges of this task for CycloCross. First, EpiLC of mouse is not a direct analog of iMeLC of human. Then, note that human-mouse divergence is greater than human-macaque meaning inference of human states from mouse cells is more complex. This is confirmed by the performance of using the expression value of ESC of macaque as a substitute for the expression value of ESC of human, which is a better approximation than using the expression value of EpiLC of mouse to replace the expression value of iMeLC of human.

### CycloCross generates realistic scRNA-seq data

By symmetrically translating scRNA-seq data between two species, CycloCross generates in silico data for each species. We evaluate CycloCross performance by generating human cells from the PGCLC induction process using two source datasets: macaque cells collected during the PGCLC induction process (8,432 cells, 1,180 transcription factors) and mouse cells collected during the PGCLC induction process (1,761 cells, 786 transcription factors). Data description in the Methods section provides a detailed description of the datasets. We benchmarked CycloCross against scGAN, a Wasserstein GAN designed for single-cell RNA-seq data generation [34], implemented with its default parameters and architectures as provided in their public repository [33]. In preliminary experiments, we trained a Wasserstein GAN using CycloCross’s generator and discriminator architectures, and conversely, trained CycloCross with scGAN’s default architectures. Both configurations yielded poor performance with highly unrealistic cell samples, reflecting the well-documented difficulty in GAN training [3].

We visualize the complete set of 11,277 human cells and 1,000 simulated cells, generated by sampling points in the latent space for scGAN and by randomly sampling macaque or mouse cells for CycloCross (Fig. 3a). Mode collapse, a prevalent GAN failure mode in which generated samples inadequately represent input data diversity [26], is visually absent in CycloCross results; the simulated cells show a distribution qualitatively consistent with true cell populations. In contrast, scGAN-generated cells fail to fully overlap with true cells, indicating both mode collapse and synthetic cell artifact. To quantitatively assess generation quality, we employed a rigorous evaluation framework (Figs. 3b and 3c). We withheld 1,000 cells as a test set and trained both methods on the remaining human data. CycloCross additionally utilized all available macaque and mouse cells. After training, we generated 1,000 cells for each method (via latent space sampling for scGAN and cross-species translation for CycloCross). Three complementary metrics were computed: Euclidean distance between real and generated cells, cosine distance, and random forest classification accuracy. For the classification metric, we first reduced dimensionality to 50 principal components following the GroundGAN pipeline [61], then trained a random forest with a 70-30 train-test split. We compute the area under the ROC curve (AUROC) (Fig. 3b). A perfect generative method achieves AUROC = 0.5, as classifiers should be unable to distinguish real from synthetic data. Both methods are compared with a "control sample", where the test cells are compared with 1,000 cells randomly picked from the training set. The values for the control sample give a lower bound in terms of the accuracy of any method. Results were averaged across five random train-test splits and five independent cell generations of 1,000 cells each (detailed evaluation protocol in Methods).

**Figure 3:**
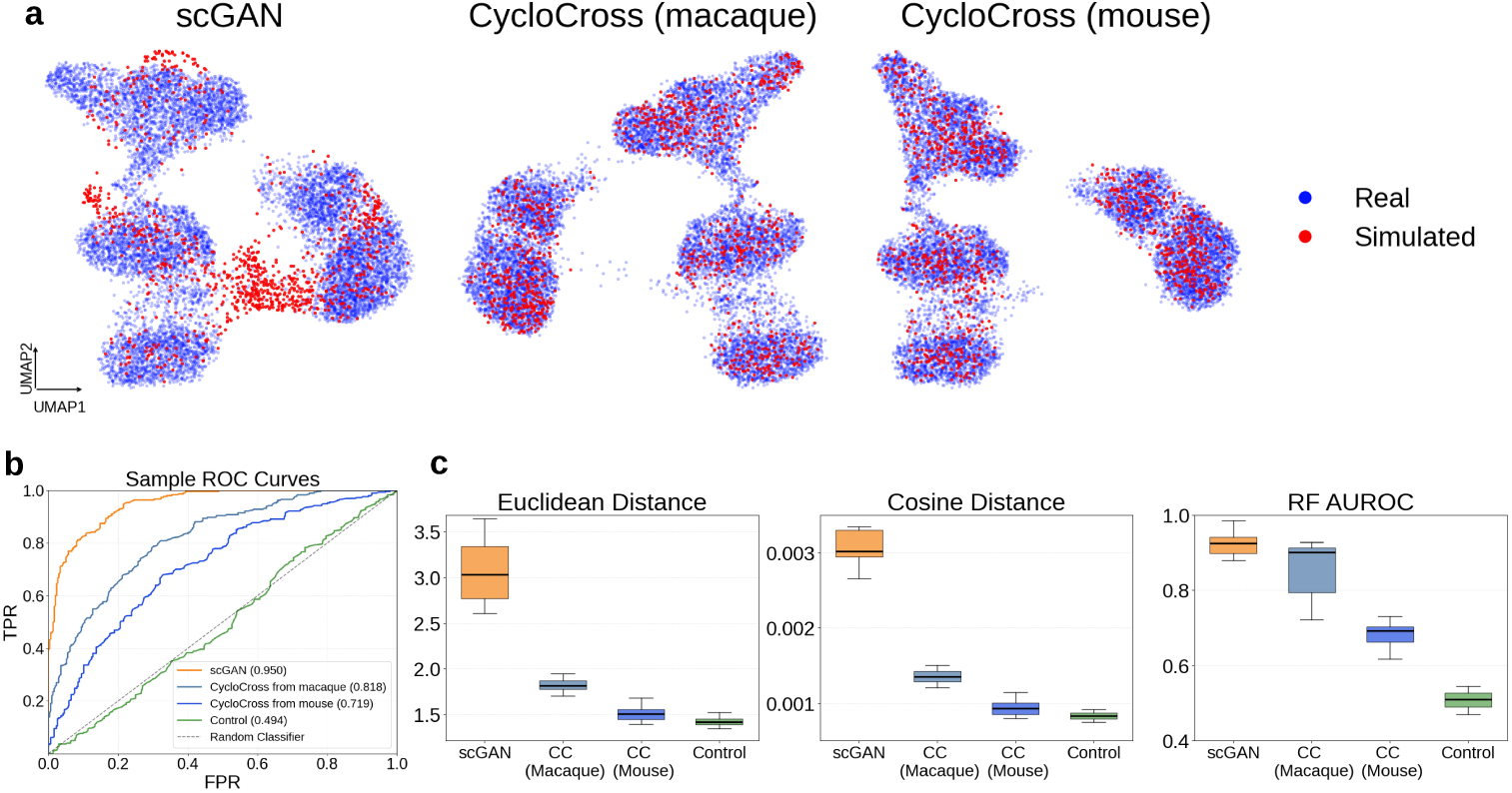
Evaluation of CycloCross for the data generation task. **a**, UMAP plot of real and simulated data for scGAN, and mouse and macaque data translated from CycloCross. **b**, The ROC curve of a Random Forest classifier in distinguishing between simulated and real cells. AUROC values in brackets in the legend. **c**, Box plots of CycloCross (CC) and scGAN for human PGCLC data generation across 3 metrics: Cosine distance, Euclidean distance and AUROC.

Here, we can see that CycloCross, by using data from a related species as a starting point, achieves a much better accuracy than a WGAN-based method. In particular, the performance in terms of Euclidean and Cosine distance is very strong (and close to the control value for translated mouse data), indicating a faithful spatial organization of the generated cells. In terms of realism, cells generated by CycloCross also tend to fool a random forest more often than scGAN. By being specifically adapted to tackle inter-species translation, it is natural that CycloCross outperforms a simple noise-to-data generator. Unfortunately, it is currently impossible to compare CycloCross to other cross-species methods. Indeed, methods like SATURN [44] and CONCORD [60] align data across species but currently do not provide a pipeline to translate unobserved data from a species into data from another species. We further attempted to evaluate our method against Icebear [55] using their publicly available implementation [54]. However, the code contained unresolved errors that prevented successful execution, and we could not replicate their reported results.

A notable finding emerged: mouse-translated cells consistently outperformed macaque-translated cells across all three metrics, despite the closer phylogenetic proximity between humans and macaques. This result may reflect the extensive knowledge accumulated from mouse and human studies, together with mature gene annotations and experimental resources in both species, which may make mouse data analytically more compatible with human data. This observation aligns with recent integration benchmarking studies demonstrating comparable performance between human-macaque and human-mouse data integration for some batch correction methods, see [57, Figure 4].

**Figure 4:**
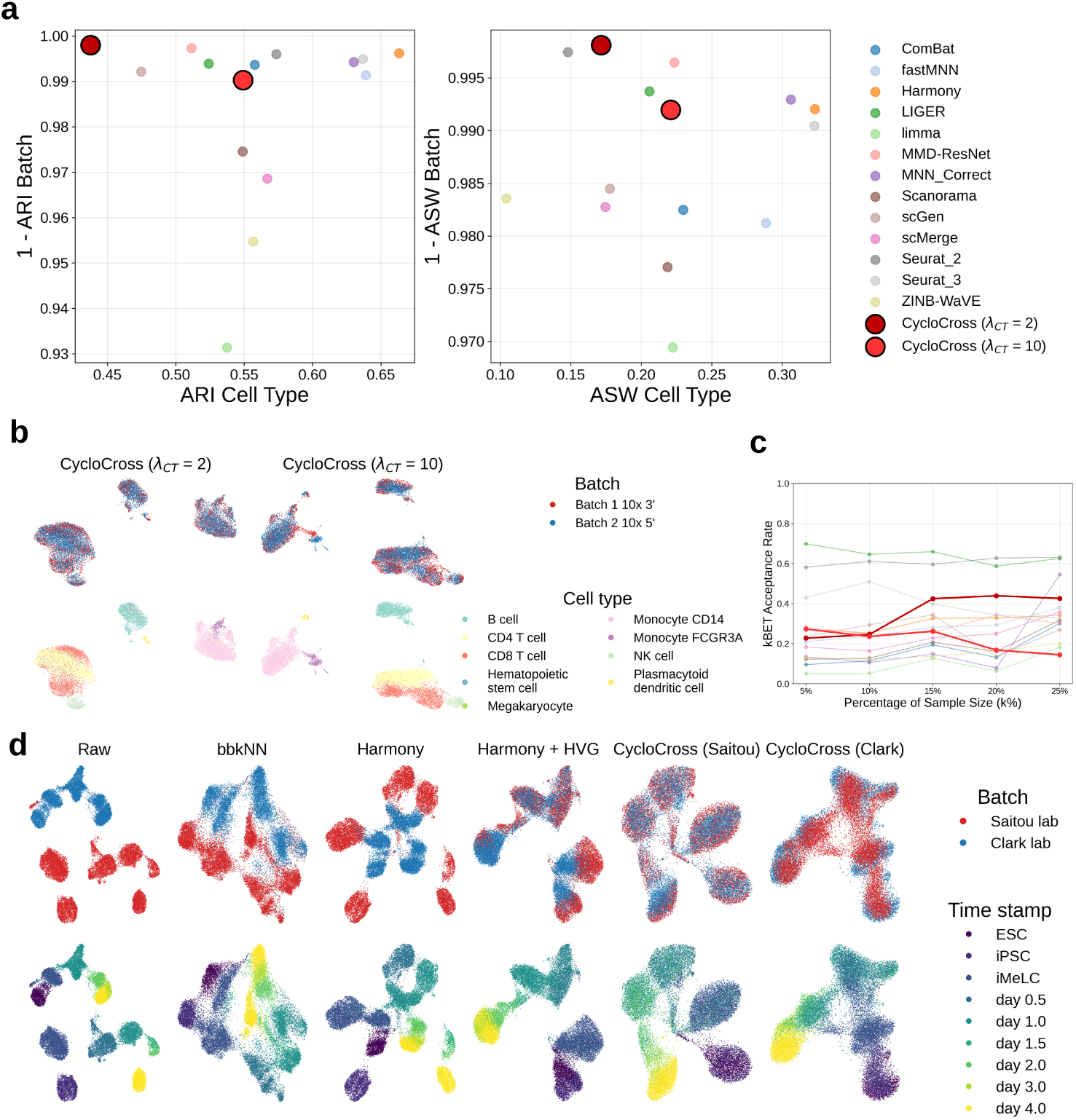
Comparison of CycloCross with other batch integration methods. **a**, ASW and ARI metric for CycloCross and 13 other batch integration methods for batch and cell type clustering for the human PBMC dataset. The best methods lie in the upper-right part. **b**, UMAP plot of the results of CycloCross for the PBMC dataset, colored by batch and cell type. **c**, Acceptance rates for kBET over a range of different *k* for the blood cells dataset for CycloCross and 13 other batch integration methods. High values denote good batch mixing. **d**, UMAP plots of the results of CycloCross and other batch integration methods for the high-dimensional PGCLC data set, colored by batch and time stamp

Finally, note that a specificity of CycloCross is that it can only generate as many scRNA-seq data from *S*_2_ as we have available cells of *S*_1_. If we want to generate a large number of cells of *S*_2_, we should first use a generative method based on a latent space sampling, such as GANs [34], variational auto-encoders [13] or diffusion models [28] to simulate cells of *S*_1_. CycloCross can then be used to translate these simulated cells into simulated cells of *S*_2_.

### CycloCross removes batch effects through same-species translation

Although we have so far focused on cross-species translation, CycloCross can also be applied to datasets obtained from the same species. In this setting, the method maps one dataset into the expression space of another dataset from the same species. When the two datasets contain comparable cell populations or cell states, this translation can be interpreted as removing dataset-specific batch effects while preserving the underlying biological structure.

We assess the capacity of CycloCross to translate data from the same species acquired in different experimental conditions such as different protocols or labs. Merging data from different experimental setups is a primary source of batch effects [11]. There has been extensive works to mitigate these effects a some review work to compare these methods in detail [50]. We here illustrate the capacity of CycloCross to remove batch effects.

We first validate our approach using a benchmark human peripheral blood mononuclear cell (PBMC) dataset [56], combining two batches generated with the 10x Genomics 3’ and 5’ protocols. Since both datasets were generated from PBMCs derived from the same donor using closely related protocols, they are expected to contain relatively modest batch effects compared with datasets generated across different laboratories or biological sources. The data were preprocessed to include 8,098 cells (first batch) and 7,378 cells (second batch), spanning 17,430 genes [50]. The dataset comprises nine distinct cell types.

Before applying CycloCross, we selected the 1,000 most variable genes using Seurat v3 [46]. CycloCross was conditioned on each cell type using one-hot encoding. We evaluate its ability to integrate batches while preserving cell-type information. We compute the Average Silhouette Width (ASW) and Adjusted Rand Index (ARI) between *k*-means clustering results and ground truth labels for both cell type and batch. An optimal method should yield ASW and ARI values close to 0 for batch prediction (indicating indistinguishable batches) and close to 1 for cell type prediction (indicating preserved biological structure). We compare the results obtained for two values of the batch-to-cell trade-off parameter *λ*_CT_ against 13 other integration methods and raw input (Fig. 4a). Scores for competing methods were extracted from the benchmark paper [50]. Additionally, we assess batch integration using the *k*-nearest neighbor Batch Effect Test (kBET) [6]. For each cell, we compare the local batch distribution among its *k* nearest neighbors to the global batch distribution using a *χ*^2^ test. Acceptance rates are calculated for *k* values of 5%, 10%, 15%, 20%, and 25% of the total sample size (Fig. 4c). Here again, CycloCross is benchmarked against 13 other integration methods. High kBET acceptance rates across *k* values indicate effective batch mixing. We refer to Methods for the precise evaluation pipeline.

We report the results for the integration in the space of the first batch. We plot the UMAP of the integrated cells with CycloCross (Fig. 4b). The results for the other 13 methods are present in [57, Figure 4]. For both values *λ*_CT=2_ and *λ*_CT_ = 10, the batch integration is excellent, while the cell types are well preserved. In agreement with the computations of ARI and ASW, taking a higher value for *λ*_CT_ enables a better cell type separation. In particular, setting *λ*_CT_ = 10 allows to separate the CD4 and CD8 T cells, while they appear to be mixed when setting *λ*_CT_ = 2. For a small *λ*_CT_ (*λ*_CT_ = 2), CycloCross achieves state-of-the-art batch integration, rendering batches indistinguishable in k-means clustering (Fig. 4a). The kBET scores (Fig. 4c) confirm excellent local-to-global batch distribution alignment, outperforming most competing methods. However, small *λ*_CT_ values compromise cell-type identification. Increasing *λ*_CT_ to 10 balances both objectives, maintaining top-tier performance in batch integration and cell-type preservation, with acceptable kBET mixing scores.

We next evaluate CycloCross in a more challenging batch-integration setting using human cells from the PGCLC induction process starting from induced pluripotent stem cells (iPSCs). In contrast to the PBMC benchmark, these datasets were obtained from distinct individuals and laboratories, and were collected at different time points and stem cell states, resulting in substantially stronger biological and technical heterogeneity. The two datasets comprise 15,203 and 11,277 cells, respectively; detailed information on the datasets is provided in the Methods section.

We apply CycloCross with *λ*_CT_ = 5 to these datasets. As a preprocessing step, we restricted the analysis to approximately 1,000 transcription factor genes. For comparison, bbkNN [42] and Harmony [25] were applied after reducing the data to 50 principal components, following the standard usage recommended for these methods. We compare the alignments of the two batches by displaying UMAP plots of the integrated datasets (Fig. 4c).

Although bbkNN and Harmony partially connected clusters corresponding to some matched time points, they did not achieve cell-level integration across the two datasets. When Harmony was applied after selecting 200 highly variable genes followed by 50-principal components, the integration improved for several time-point clusters, but the two datasets were still not globally merged. These results suggest that the batch effects between the two laboratories were substantially larger than those typically assumed by standard batch-correction methods, and that the magnitude of the batch effect may vary across time-point-specific clusters.

In contrast, CycloCross successfully merged the two datasets across all time-point clusters without requiring PCA or other low-dimensional embeddings. Moreover, CycloCross recovered the temporal continuity of the PGCLC induction process after integration. This improvement is likely because CycloCross does not simply merge similar clusters under a predefined similarity criterion, as in conventional batch-correction approaches. Instead, it learns an explicit translation from one dataset to the other, which may allow it to overcome both large batch effects and cluster-dependent heterogeneity in batch noise. This stronger alignment is supported quantitatively by the ARI and ASW metrics for batch and time-stamp clustering in Table 1.

**Table 1:**
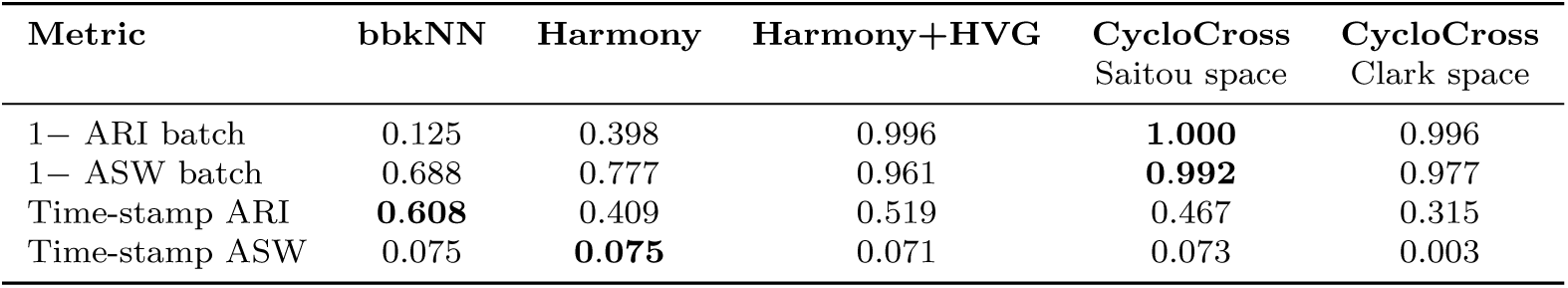
Comparison of bbkNN, Harmony and CycloCross for batch integration of human PGCLC datasets.

| Metric | bbkNN | Harmony | Harmony+HVG | CycloCross<br>Saitou space | CycloCross<br>Clark space |
| --- | --- | --- | --- | --- | --- |
| 1- ARI batch | 0.125 | 0.398 | 0.996 | **1.000** | 0.996 |
| 1- ASW batch | 0.688 | 0.777 | 0.961 | **0.992** | 0.977 |
| Time-stamp ARI | **0.608** | 0.409 | 0.519 | 0.467 | 0.315 |
| Time-stamp ASW | 0.075 | **0.075** | 0.071 | 0.073 | 0.003 |

Another important advantage of CycloCross is that it does not require the two datasets to be defined on the same gene set. In contrast, conventional batch-integration methods such as bbkNN and Harmony require a shared feature space; therefore, we restricted the comparison to 966 transcription factors common to both datasets when applying these methods. CycloCross can instead use the original gene sets of each dataset, thereby avoiding the loss of information caused by restricting the analysis to shared genes.

A consequence of this formulation is that CycloCross integrates two datasets by translating one dataset into the expression space of the other, rather than embedding both datasets into a common latent space. Thus, either dataset can be used as the reference space: the Saitou lab data can be translated into the Clark lab space, or the Clark lab data can be translated into the Saitou lab space. This procedure is not symmetric, and the choice of reference dataset can affect the integration quality. Indeed, Table 1 shows that using the Saitou lab data as the reference space yields slightly better preservation of time-stamp structure.

We have demonstrated that, both qualitatively and quantitatively, CycloCross is a powerful batch integration method that can integrate data from different sources while preserving the organization of the time stamps or cell type information.

### Identification of contrastive genes between species

In addition to its generative properties, CycloCross can be used as an analysis tool to examine differences between species. In particular, CycloCross enables the identification of genes that exhibit contrastive expression patterns between *S*_1_ and *S*_2_. Here, the translation of gene expression between species remains a biological and computational challenge, because orthologs do not always preserve equivalent regulatory roles, expression patterns, or cell-type-specific functions. Identifying such contrastive genes allows us to further characterize evolutionary differences between species. This approach is complementary to that of Species-OT [49] which infers cross-species gene correspondences by focusing on the comparable geometric structure of transcriptome data arising from well-conserved underlying gene regulatory networks.

We aim to discriminate between real data from *S*_1_ and translated data generated from data from *S*_2_. We formulate this as a binary classification problem with samples (*z_i_, r_i_*) (*i* = 1*, . . ., N*_1_ + *N*_2_), where *z_i_ ∈* R*^p^*^1^ is either a real cell from *S*_1_ or a translated *S*_1_-like cell generated from *S*_2_, and *r_i_* = 1 for true cells, *z_i_ ∈ {X_ℓ_}*, and *r_i_* = 0 for translated cells, *z_i_ ∈ {G*_21_(*Y_ℓ_*)*}*. We train a ridge-regularized linear classifier [17] to solve this problem. For each cell *z*, the classifier predicts a label *ri*^∽^ defined as

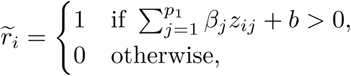

where *z_ij_* denotes the expression value of gene *j* in the cell *z_i_*, and the coefficients *β_j_* and the intercept *b* are learned so that the predicted labels *r_i_*^∽^ match the true labels *r_i_* as accurately as possible under ridge regularization.

We define the relative gene importance *γ_j_* of gene *j* as

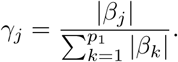

The gene importance corresponds to the contrastive power of gene *j* to distinguish real from translated data. A high value means this gene is, relatively to others, preferred for classification and therefore that its estimation from genes of the other species is difficult for CycloCross. This suggests that this gene could be a marker of inter-species differences (Fig. 5a).

**Figure 5:**
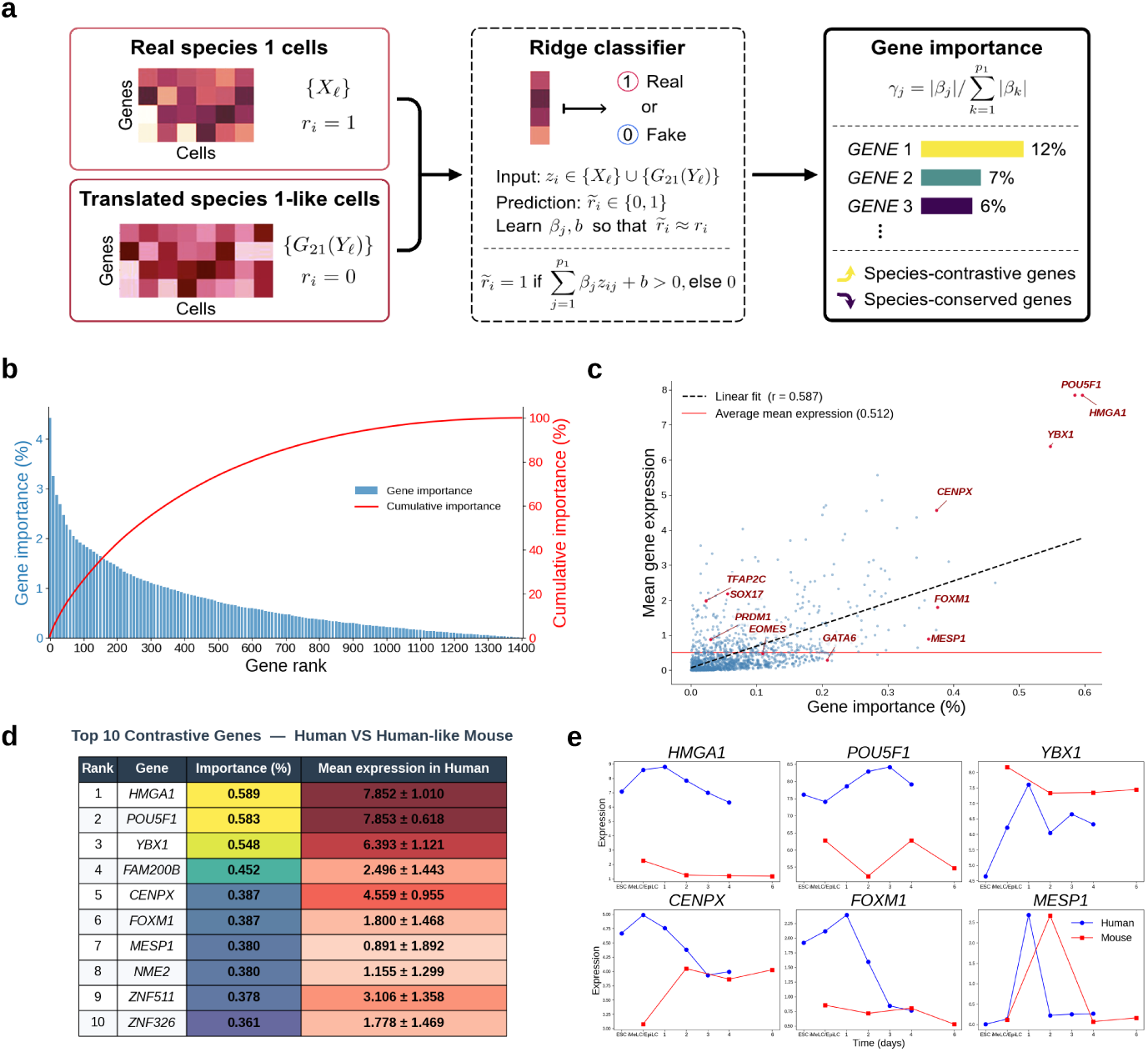
Identification of most contrastive genes. **a**, Global pipeline. We train a classifier to distinguish between real and translated data. The top *k* genes with the highest relative importance for the classification task are identified as *contrastive genes*. **b** Evolution of the gene importance according to its rank when classifying human from human-like mouse data. Each bar corresponds to 10 genes. **c** Mean gene expression as a function of gene importance for each human gene used in the study. **d**, List of the 10 most contrastive genes when classifying human data and human-like mouse data. Gene importance and mean expression level with the standard deviation of each gene are provided. **e** Expression evolution of the six most contrastive genes present in both human and mouse.

We apply CycloCross to human and mouse PGCLC induction datasets. PGCLC induction provides a biologically well-characterized setting for cross-species comparison because it models the emergence of the germline lineage, a process that is broadly conserved across mammals but controlled by partially species-specific regulatory programs [23, 24]. Therefore, genes that distinguish real human PGCLC trajectories from human-like mouse trajectories may highlight regulatory components that are not fully recapitulated by the mouse-to-human translation.

We remark that there is a sharp initial decay in the importance of the genes (Fig. 5b) with about 10 % of the total importance concentrated in the first 30 genes. We also demonstrate that the gene importance is positively correlated to the average gene expression (Fig.5c). In particular, the three most contrastive genes *HMGA1*, *POU5F1* and *YBX1* are also the three genes with the highest level of expression on average. However, this correlation is somehow loose, with a correlation coefficient *r* = 0.588. In practice, it means that some genes, such as *TFAP2C* can have high expression levels without being contrastive.

We selected the ten genes with the highest gene importance *γ_i_* for distinguishing real human data from human-like mouse data (Fig. 5d). These highly contrastive genes showed clear differences between human and mouse in their expression levels and/or temporal expression patterns (Fig. 5e). For example, *HMGA1* and *POU5F1* showed markedly different expression levels, whereas *YBX1*, *CENPX*, and *FOXM1* showed different temporal patterns, and *MESP1* showed a delayed pattern. Inter-species differences in developmental timing are fundamental. Here, CycloCross identifies contrastive genes that exhibit similar expression patterns with temporal offsets, advancing the characterization of species-specific biological clocks. We show the expression of a few highly contrastive genes that shows a delayed pattern between human and mouse in Figure S2 of the supplementary material.

Interestingly, several canonical regulators of human PGCLC specification were not highly ranked in this contrastive analysis, even though they showed relatively high expression levels in the human data. For example, the ranks of *EOMES*, *SOX17*, *TFAP2C*, and *PRDM1* were 317, 610, 1,229, and 789, respectively. This may be because functionally corresponding regulatory factors are also present in the mouse PGCLC program, including *T*, *TFAP2C*, *PRDM1*, and *PRDM14*, which may partially compensate for species-specific differences in the translation task. In contrast, *MESP1* and *GATA6*, which have recently been identified analytically as potential inhibitors of human PGCLC differentiation [53], were ranked 8th and 76th, respectively. This suggests that these genes may represent human-specific barriers to PGCLC differentiation that are not fully captured by the mouse-derived translated profiles.

Some of the top-ranked genes have documented human–mouse divergence. Most notably, in human PGCLCs, *POU5F1* partners with *SOX17* to drive germline identity, whereas in mouse PGCLCs it partners with *SOX2*, indicating that the same pluripotency factor can participate in distinct transcriptional complexes depending on the species [9, 47]. Other examples include *HMGA1*, which has additional exons in its 5’ untranslated region in humans but not in mice, suggesting species-specific transcriptional regulation [41], and *FOXM1*, which is required for embryonic stem cell pluripotency in mice, whereas its role in human embryonic stem cells is less clear and may involve different regulatory mechanisms [51]. Conversely, some top-ranked genes, such as *YBX1* [1] and *CENPX* [35], are relatively conserved between human and mouse. These cases indicate that high contrastive importance may reflect not only biological divergence but also numerical discrepancies or limitations of the translation model. Finally, genes with less well-characterized functions, such as *FAM200B* and *ZNF511*, may provide candidates for future investigation of species-specific regulation during PGCLC induction.

## Discussion

In this study, we present CycloCross, a versatile translation pipeline designed to bridge single-cell gene expression data across species. By leveraging a machine learning framework with adversarial training, CycloCross enables the transformation of cells from one species into another, ensuring that the translated cells retain biological realism and fidelity. Our extensive evaluations on primordial germ cell-like cell (PGCLC) induction systems from human, macaque and mouse demonstrate that CycloCross excels in correcting batch effects between data samples from the same species, generating simulated cells based on real cells from a different species, and preserving the time evolution of data when translating cells between species.

CycloCross is not restricted to cross-species applications; it can translate data between any two distinct sources, effectively functioning as a batch correction method where cells from different datasets are harmonized into a shared latent space. This enables joint analysis of translated and original cells. Benchmarking against state-of-the-art batch correction methods reveals that CycloCross consistently ranks among the top performers, offering a tunable trade-off between batch indistinguishability and preservation of cell-type organization. Its heavy architecture allows for exceptional batch correction when appropriately tuned, making it particularly effective for complex datasets with significant batch effects. However, its sole use as a batch integration method must be mitigated. Indeed, it is a heavy method relying on complex mappings using large neural networks between two gene spaces. This means that the computation cost is greatly increased as opposed to more standard methods. Integrating two batches of approximately 10,000 cells each and 1,000 genes takes about an hour for CycloCross while it is in the order of minutes for Harmony and bbkNN. Furthermore, the complexity of the mapping obtained makes it less interpretable than more simple methods. Finally, we remark that although CycloCross could be easily generalized to the integration of *B* batches with *B >* 2, this would require training 2(*B −* 1) generators and *B* discriminators, which is a strong computational caveat as opposed to methods such as Harmony that are naturally designed to handle multi-batch data.

CycloCross is particularly well-suited for handling highly dissimilar datasets, such as those encountered in multi-species studies. It can manage datasets with completely different gene sets, a challenge that often limits the applicability of traditional methods. In comparative analyses, CycloCross-generated data were found to be significantly more realistic than those produced by standard Wasserstein GANs. This is natural: as a GAN learns how to generate a realistic sample from a random vector, CycloCross starts from a real cell from a different species. Thanks to the evolutionary similarities between species, the mapping that CycloCross has to learn is much simpler, which translates in an improved performance. In the future, we would like to integrate more species in the analysis. We demonstrated that translated mouse cells were more realistic human cells than translated macaque cells. This seemingly surprising fact should be investigated further, by including more distant species and verifying if there is an overall performance decay consistent with findings from recent benchmarking studies [57]. This finding is especially promising for real-world applications. Since mouse data is far easier and more cost-effective to acquire than macaque data, the ability of CycloCross to maintain high realism even across more distant species opens exciting new possibilities. Expanding the pool of viable experimental species could dramatically reduce barriers to research and accelerate scientific progress.

CycloCross can also be used as a tool to analyze the data. Once trained, it can be used to analyze genes that display strong inter-species divergence. The rationale is that if a linear classifier manages to separate the real data from the translated data using some key set of genes, these are non accurately translated by CycloCross. As we demonstrated that CycloCross generates realistic samples across a wide range of metrics, this points out towards these genes as being potential markers of inter-species divergence. Notably, several of the identified candidates remain poorly annotated or functionally uncharacterized, highlighting potential targets for downstream experimental validation. This approach not only prioritizes genes for evolutionary studies but also provides a computationally efficient, hypothesis-generating framework to guide wet-lab or comparative genomics research. Furthermore, we saw that CycloCross could successfully identify genes with a delayed expression between species. For PG-CLC induction data, there is only a small delay of at most a couple of days between species. Applying this analysis to slowly developing organs, such as the brain, could identify genes with delayed expression and quantify inter-species temporal offsets in days, providing new insights into developmental differences.

The primary contribution of this work lies in the application of CycloCross to PG-CLC and stem cell data, where we demonstrate its ability to efficiently translate cells between species. In particular, when a cellular state is missing for a target species but known for a source species, CycloCross can translate this state with high precision. This capability is crucial, as it allows researchers to bypass ethical, financial, or practical constraints associated with acquiring specific data for certain species. CycloCross could thus be applied to a wide range of scenarios, including translating disease progression trajectories or treatment response profiles from model organisms to humans. Additionally, its capacity to predict the outcomes of gene knock-out experiments—by leveraging real data from a source species and wild-type data from a target species—highlights its potential to revolutionize experimental design and hypothesis testing in systems biology.

While this study focuses on single-cell RNA sequencing (scRNA-seq) data, the adversarial architecture of CycloCross is inherently generalizable to different data modalities. Indeed, the original CycleGAN paper [59] was originally defined for images. Adapting the architecture of the generators and the discriminators could allow to translate other different types of data. While a similar architecture using fully-connected layers could allow to handle scATAC-seq or scDNA-seq data, including convolution layers could allow to handle image data, and attention heads would allow to translate epigenomics and chromatin contact data [10].

Finally, note that in this paper, except when evaluating the capability of CycloCross to generate realistic samples, the entire set of available cells was used for the data translation. The optimization of the weights of the networks in CycloCross was retrained from scratch when applied to a new pair of species. Investigating the ability of CycloCross to generalize a pre-trained network to new data, possibly with a fine-tuning step, would be an important direction of research. For instance, CycloCross could be trained between two different time stamps of the same species to predict the mechanism ruling time evolution of data. It could then be applied to evaluate the effects of gene knock-out experiments. Such advancements could further enhance the scalability and efficiency of CycloCross, making it an even more indispensable tool for the single-cell genomics community.

## Methods

### Data and preprocessing

#### Data description

We consider a human peripheral blood mononuclear cells dataset [56] to validate the batch integration study. The 3’ and 5’ 10x Genomics protocols which capture different regions of mRNA were used to generate the two data batches. Preprocessed data are available online [27]. The resulting dataset consists of 8,098 cells for the 3’ batch and 7,378 cells for the 5’ batch.

For the batch integration of human PGCLC induction systems, we use two datasets. The first one, referred to as the one coming from the Saitou lab consists of 15,203 cells after preprocessing [53]. The second one, referred to as the one coming from the Clark lab consists of 11,277 cells after preprocessing [7].

For the translation tasks, we use PGCLC induction systems from the Clark lab for human and macaque and mouse data from the Saitou lab. We refer to Table 2 for a breakdown of the number of genes and the number of cells per time stamp for each species, after data preprocessing.

**Table 2:** Number of genes and cells in scRNA-seq data of PGCLC induction systems used for inter-spceies translation.

|  | Human | Macaque | Mouse |
| --- | --- | --- | --- |
| #genes | 1,404 | 1,180 | 786 |
| #cells |  |  |  |
| ESC | 1,878 | 2,116 | NA |
| EpiLC | NA | NA | 1,236 |
| iMeLC | 2,108 | NA | NA |
| Day 1 | 3,282 | 2,758 | NA |
| Day 2 | 1,490 | 246 | 167 |
| Day 3 | 1,116 | NA | NA |
| Day 4 | 1,403 | 1,317 | 336 |
| Day 6 | NA | 1,035 | 22 |
| Day 8 | NA | 960 | NA |
| Total #cells | 11,277 | 8,432 | 1,761 |

#### Preprocessing

The PGCLC dataset has been preprocessed according to the following pipeline. First, we filtered cells to ensure the homogeneity of the cell population. At each time point, we selected cells with high expression of both *PRDM1* and *TFAP2C*. To do this, we first applied the denoising algorithm RECODE [20, 21], performed a log-normalization, and generated a UMAP embedding. We then manually selected cells from clusters characterized by high expression of both *PRDM1* and *TFAP2C*. We then removed cells with fewer than 100 genes and genes expressed in less than 5 cells. We then applied RECODE again to mitigate the impact of acquisition noise in high-dimensional data. To adjust for sequencing depth and ensure accurate comparison of gene expression levels, we normalized the total read count to 10^5^ so that every cell has the same total count [19]. We then log-normalized the expression counts via *x →* log_2_(1 + *x*) [30, 62]. Finally, we filtered the set of genes by keeping only transcription factor genes, which regulate gene expression and constitute a core basis of the entire gene set. A list of human transcription factor genes is available [29], and we can also use it to slice data of other species based on the gene names, assuming well-conservation of transcription factors across closely related mammalian species. This significantly reduce the dimension of the data.

### Optimization details

We present the architecture and hyperparameters of the neural networks used to train CycloCross. These choices were manually calibrated using the batch integration task on PGCLC data from the Saitou and Clark labs. Our quality criterion was to select parameters that minimize the AUROC of a random forest classifier trained to distinguish between real data from the Saitou lab, and Saitou lab-like data translated from the Clark lab.

#### Networks architectures

CycloCross architecture relies on four neural networks:

- Two generators that each take a cell from a species and transform it into the cell of another species. These generators are fully connected networks constituted of four hidden layers of 1024-2048-4096-4096 neurons. We consider Leaky ReLU activation functions and batch normalization between each layer.
- Two discriminators that are binary classifiers neural networks that take a cell and return a continuous value between 0 and 1. The discriminators aim at giving the value 0 to cells simulated by the generators, and 1 to cells from the dataset. These discriminators are fully connected networks constituted of three hidden layers of 4096-2048-1024 neurons. We consider Leaky ReLU activation functions and batch normalization between each layer.

We optimize the CycleGAN loss

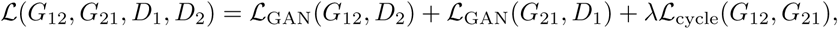

using the ADAM optimizer [22]. To improve stability, we penalize the norm of the gradient of the discriminator [14].

#### Hyperparameters

The training of CycloCross involves some hyperparameters. We refer to Table 3 for a detailed description of each parameter and the default value.

**Table 3:** Description of hyperparameters used by CycloCross.

| Parameter | Description | Default value |
| --- | --- | --- |
| $N_{\text{it}}$ | Number of iterations of the gradient descent | 16000 |
| LR | Learning rate | Decreasing between 0.00002 and 0.00001 |
| $\lambda$ | Trade-off between GAN loss and cycle loss | 1 |
| $\lambda_{\text{CT}}$ | Trade-off between gene expression and cell-type | 5 |
| $\lambda_{\text{GP}}$ | Gradient penalty | 4 |
| $\vert B\vert $ | Batch size | 512 |
| $G : D$ | Number of generator steps per discriminator step | 5 |

### Evaluation methods

#### Pseudo-bulk gene expression interpolation

For each species *S*_1_ and *S*_2_, assume we dispose of time stamps 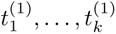 and 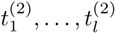 . Note that the time stamps are not necessarily matching between each species. For each gene *g*, we average the gene expression for each time stamp in order to obtain the time series: 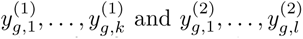. Assume that a time stamp *t_⋆_* is present only for *S*_2_ and not for *S*_1_. We can then translate every cell at *t_⋆_* for *S*_2_ into cells of *S*_1_ using CycloCross. We average the expression of these translated cells at time stamp *t_⋆_* to obtain an estimate of the expression value 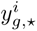 at time *t^⋆^*.

We compare CycloCross to two other approaches:

- Using the value 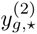 known for *S* to approximate 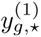
- Fitting a cubic spline to the known data 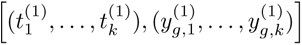 and then evaluating it at *t⋆*.

For every method, the performance of the predictor *y*^ is measured by averaging over the set *O* = *{*genes present in both *S*_1_ and *S*_2_*}*:

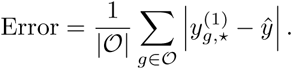

#### Resemblance of real and simulated cells

We use three metrics to evaluate the resemblance between real and simulated cells. We reproduce the pipeline of GroundGAN [61] for the sake of consistency with the literature. We first compute the centroids of real cells and simulated cells, where the centroid was obtained by calculating the mean along the gene axis. We then compute the Euclidean and cosine distance between the centroids. A low Euclidean or cosine distance means simulated cells mix well with real cells.

For the third metric, we first reduce the data to 50 principal components. We then perform a 70-30 random train-test split. We consider the binary classification problem of distinguishing real cells from simulated cells. We then train a random forest classifier with 1000 trees and computer the Area Under its Receiver Operating Characteristic (AUROC) curve. If the real cells are indistinguishable from the simulated cells, the random forest will not do better than a random guess, which has a AUROC of 0.5. The worst possible value is 1 and corresponds to the case where the Random Forest perfectly discriminates real from simulated data.

#### Batch integration

We use three metrics to evaluate the quality of batch integration of CycloCross and other methods. We reproduce the pipeline of the benchmark paper [50] for the sake of consistency. For the first two metrics ASW and ARI, we first subsample the data to keep 80 % of the original number of cells. We then consider the first 20 principal components. The procedure is repeated and averaged 20 times.

First, the silhouette score of a cell is computed by subtracting its average distance to other members in the same cluster from its average distance to all members of the neighboring clusters, and then dividing by the larger of the two values. The resulting score ranges from -1 to 1, where a high score denotes that the data point fits well in the current cluster, while a low score denotes a poor fit. We then average the scores across all cells to obtain the Average Silhouette Width (ASW). Here, the clusters can either denote the batches or the cell types/time stamps, providing two different ASW. An ideal method has a low score for the ASW for the batches and a high score for the ASW for the cell types.

The Adjusted Random Index (ARI) [18] can also be employed to evaluate batch correction methods in terms of cell type purity and batch mixing. The ARI measures the percentage of matches between two label lists, corrected for chance. We perform a *k*-means clustering (where *k* is set to be the known number of clusters) and use the ARI to measure the cluster agreement accuracy. Here, the clusters can either denote the batches or the cell types/time stamps, providing two different ARI. An ideal method has a low score for the ARI for the batches and a high score for the ARI for the cell types.

Finally, we use the kBET score [6] to measure batch mixing at a local level. After SVD-based dimension reduction, the k-nearest neighbors around each data point are selected to compute its local batch label distribution. Then, 10% of the data points are randomly selected to test the local batch label distribution against the global distribution. The null hypothesis of all batches being well mixed is not rejected if the local distribution is sufficiently similar to the global distribution. The fraction of rejections ranges from 0 to 1. If the fraction of rejections is close to zero, this signifies that the batches are well mixed. We reduced the data to its top 20 principal components. Because the number of nearest neighbors k as input has a big impact on the results of kBET, we ran kBET using a predefined list of *k* values. Following the example in the kBET paper, we chose the *k* input value equal to 5%, 10%, 15%, 20%, and 25% of the sample size.

### Experimental model and subject details for mouse data

#### mESC cell line

Mouse BDF1-2-1 ESCs carrying the Blimp1–mVenus (BV) and Stella–ECFP (SC) transgenes were cultured as described previously [38, 39]. Briefly, mouse ESCs were maintained in N2B27 medium supplemented with PD0325901 (0.4 *µ*M; Stemgent, cat. no. 04-2006), CHIR99021 (3 *µ*M; BioVision, cat. no. 4423), and leukemia inhibitory factor (LIF; 1,000 U ml-1; Merck Millipore, cat. no. ESG1107) on plastic plates coated with poly-L-ornithine (0.01%; Sigma-Aldrich, cat. no. P3655) and laminin (100 ng ml-1; BD Biosciences, cat. no. 354232). Cells were cultured at 37°C under 5% CO2 in air.

#### Mouse PGCLC induction

Mouse EpiLC and PGCLC induction was performed as described previously [38, 39]. For EpiLC induction, mouse ESCs were plated on human plasma fibronectin-coated plates (Merck Millipore, cat. no. FC010) in N2B27 medium containing activin A (20 ng ml-1; PeproTech, cat. no. 120-14), bFGF (12 ng ml-1; Invitrogen, cat. no. 13256029), and KSR (1%; Gibco, cat. no. 10828028). After 2 days of EpiLC induction, mouse PGCLCs were induced under floating conditions in low-cell-binding U-bottom 96-well plates using Glasgow’s minimal essential medium (GMEM; Thermo Fisher Scientific, cat. no. 11710035) supplemented with KSR (15%; Gibco, cat. no. 10828028), NEAA (0.1 mM; Thermo Fisher Scientific, cat. no. 11140-050), sodium pyruvate (1 mM; Thermo Fisher Scientific, cat. no. 11360-070), *β*-mercaptoethanol (0.1 mM; Thermo Fisher Scientific, cat. no. 21985023), penicillin/streptomycin (100 U ml-1 penicillin and 0.1 mg ml-1 streptomycin; Thermo Fisher Scientific, cat. no. 15140148), and L-glutamine (2 mM; Thermo Fisher Scientific, cat. no. 25030149). The medium was further supplemented with BMP4 (500 ng ml-1; R&D Systems, cat. no. 314BP01M), LIF (1,000 U ml-1; Merck Millipore, cat. no. ESG1107), SCF (100 ng ml-1; R&D Systems, cat. no. 455MC), and EGF (50 ng ml-1; R&D Systems, cat. no. 2028EG). Day 0 cells of PGCLC induction, corresponding to EpiLCs, and day 2, day 4, and day 6 PGCLCs were dissociated and cryopreserved in STEM-CELLBANKER at -80°C until single-cell RNA-seq library preparation.

#### Cell collection

Cryopreserved day 0 PGCLC-induction cells, corresponding to EpiLCs, and day 2, day 4, and day 6 PGCLCs were thawed and suspended in FACS buffer consisting of DMEM/F12 (Gibco, cat. no. 11320082) supplemented with 0.1% BSA Fraction V (Thermo Fisher Scientific, cat. no. 15260037). After washing with FACS buffer, cells were resuspended in fresh FACS buffer and filtered through a nylon cell strainer. Single-cell populations were collected using a FACSAria III flow cytometer (BD Biosciences) based on FSC/SSC gating.

### 10X scRNA-seq data acquisition

Single-cell RNA-seq libraries were generated using the 10X Genomics Chromium Controller and Chromium Single Cell 3’ Reagent Kits v3.1 according to the manufacturer’s instructions. For mouse samples, cell suspensions were subjected to Cell Multiplexing Oligo labeling according to the manufacturer’s instructions. Reverse transcription, cDNA amplification, and sample indexing were performed using an Eppendorf Mastercycler. The final libraries were quantified using a KAPA Library Quantification Kit (KAPA Biosystems, cat. no. KK4824), and fragment size distributions were assessed using a LabChip GX DNA High Sensitivity Kit (PerkinElmer). Pooled libraries were sequenced on an Illumina NovaSeq 6000 platform using an SP or S1 100-cycle kit (Illumina).

#### Mapping of transcriptome reads and conversion to gene expression levels

Reads were aligned to the GRCm38/mm10 reference genome using Cell Ranger v6.0.1 with the 10x Genomics pre-built mouse reference without including intronic reads.

#### Multiplet identification

For 10X scRNA-seq data, low-quality cells were defined as cells with fewer than 10,000 UMIs, fewer than 2,000 detected genes, or more than 20% of reads mapped to mitochondrial genes. Low-quality cells and putative multiplets were identified using the Scrublet Python package v0.2.3. The resulting filtered data were used for downstream analyses.

## Code availability

The preprocessed data, and the codes to train CycloCross and reproduce the results from the paper are freely available [15].

## Data availability

The macaque and mouse PGCLC data are currently private and will be released upon publication in a peer-reviewed journal.

## Acknowledgments

We thank Drs. Tomoyuki Tsukiyama, Chizuru Iwatani and Hideaki Tsuchiya of the Shiga University of Medical Science for the establishment of cynomolgus monkey ESCs. This research was supported by JST Moonshot R&D Grant Number JP-MJMS2021 (to Y.H.). This research was partially supported by JST CREST Grant Number JPMJCR24Q1 (to Y.I.) and ASHBi Fusion Research Grant 2026 (to Y.T.).

## Author Contributions

O.H., Y.T. and Y.I. designed the study; O.H. conducted the research; O.H. developed the computational method; O.H. wrote the code; O.H. analyzed the data; M.N. established the mouse cell lines; S.D. and T.N. established the macaque cell lines; Y.H. supervised the study; M.S. supervised and funded the biological experiments; O.H., Y.I., Y.T. and M.N. wrote the manuscript; and everyone reviewed the manuscript.

