## Supplementary Figures for "CycloCross: Adversarial single-cell RNA-seq data translation across species"

**Olympio Hacquard et al.**

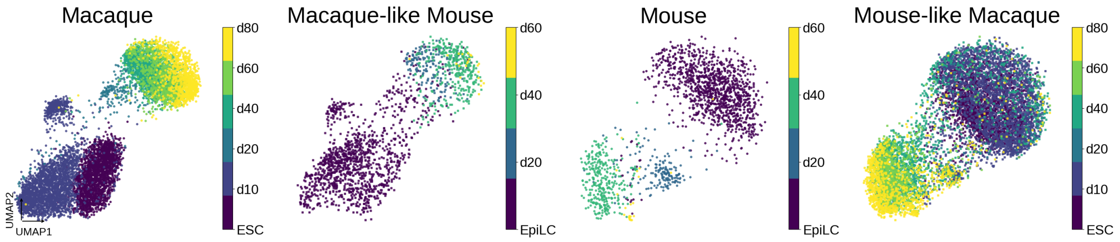


Figure S1: UMAP plots of human and macaque PGCLC induction system data translated by CycloCross, colored by time stamp. Real and simulated data are embedded in the same UMAP space, and then plotted separately.

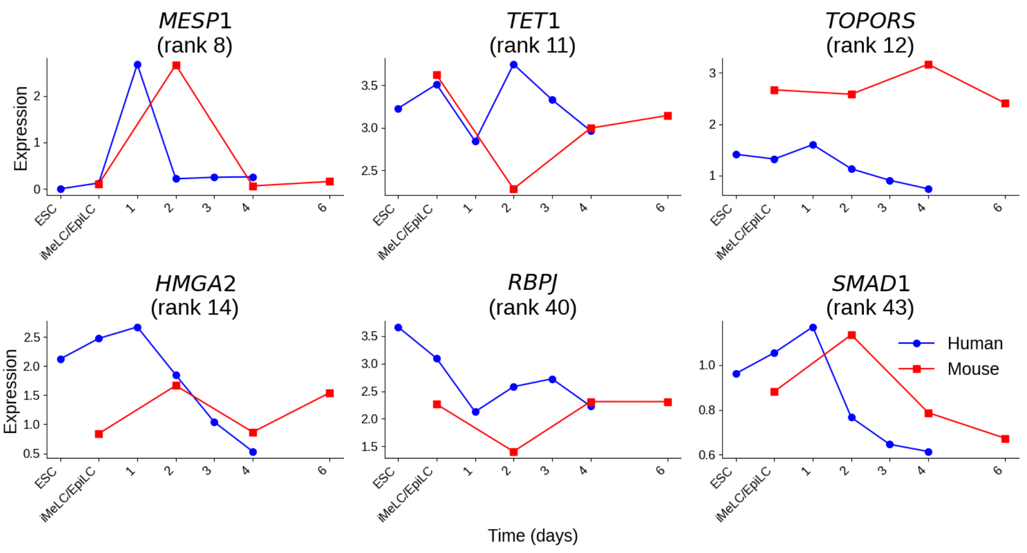


Figure S2: Highly contrastive genes with a delayed expression between human and mouse
